# A nanobody-enzyme fusion protein targeting PD-L1 and sialic acid exerts anti-tumor effects by affecting tumor associated macrophages

**DOI:** 10.1101/2024.06.05.597674

**Authors:** Yongliang Tong, Runqiu Chen, Xinrong Lu, Cuiying Chen, Guiqin Sun, Xiaolu Yu, Shaoxian Lyu, Meiqing Feng, Yiru Long, Likun Gong, Li Chen

**Affiliations:** Department. of Medical Microbiology, Key Laboratory of Medical Molecular Virology of Ministries of Education and Health, School of Basic Medical Sciences, Fudan University, Shanghai, China; Shanghai Institute of Infectious Diseases and Biosecurity, Fudan University, Shanghai, China; Department of Microbiological and Biochemical Pharmacy, School of Pharmacy, Fudan University, Shanghai, China; State Key Laboratory of Drug Research, Shanghai Institute of Materia Medica, Chinese Academy of Sciences, Shanghai, China; Department of Research and Development, Sysdiagno (Nanjing) Biotech Co., Ltd, Nanjing, Jiangsu Province, China; School of Medical Technology and Information Engineering, Zhejiang Chinese Medical University, Hangzhou, Zhejiang Province, China; Translational glycomics research center, Fudan Zhangjiang Institute, Shanghai, China

## Abstract

Cancer cells employ various mechanisms to evade immune surveillance. Their surface features, including a protective “sugar coat” and immune checkpoints like PD-L1 (programmed death ligand 1), can impede immune cell recognition. Sialic acids, which carry negative charges, may hinder cell contact through electrostatic repulsion, while PD-L1 transmits immunosuppressive signals to T cells. Furthermore, cancer cells manipulate macrophages within the tumor microenvironment to facilitate immune escape. Prior research has demonstrated the effectiveness of separately blocking the PD-L1 and sialic acid pathways in eliciting anti-tumor effects. In this study, we investigated the relationship between PD-L1 expression and genes associated with sialic acid in clinical databases. Subsequently, we developed a novel nanobody enzyme fusion protein termed Nb16-Sia to simultaneously target both PD-L1 and sialic acid pathways. In vivo experiments confirmed the anti-tumor activity of Nb16-Sia and highlighted its dependence on macrophages. Further investigations revealed that Nb16-Sia could polarize macrophages towards the M1 phenotype through the C-type lectin pathway in vitro and eliminate tumor-associated macrophages in vivo. In conclusion, our findings demonstrate that the fusion of PD-L1 nanobody with sialidase effectively targets tumor-associated macrophages, resulting in significant anti-tumor effects. This approach holds promise for drug development aimed at enhancing immune responses against cancer.

## Introduction

Antibody fusion proteins represent a category of antibody-based constructs that tether diverse payloads, including cytokines, toxins, enzymes, neuroprotectants, and soluble cytokines, to various antibody components such as full-length antibodies, Fc domains, single-chain variable fragments (scFvs), single-domain antibodies (nanobodies), and antigen-binding fragments (Fabs)[1, 2]. Several antibody fusion proteins have received regulatory approval as therapeutic agents, underscoring their significant clinical utility[3, 4]. Leveraging antibodies for targeting can markedly augment payload potency while mitigating adverse effects[5–7]. Additionally, protein payloads fused with antibodies demonstrate decreased renal clearance and prolonged in vivo half-life, thereby reducing the necessity for frequent dosing[8].

PD-L1, initially cloned in 1999, emerged subsequently as the principal ligand for PD-1 (programmed cell death 1) on T cell surfaces, modulating immunosuppressive effects[9, 10]. Elevated expression levels of PD-L1 have been observed in tumor cells and macrophages within tumor tissues, facilitating immune evasion[11, 12]. Antibodies targeting PD-L1 have proven instrumental in bolstering anti-tumor immune responses[13]. Nevertheless, notwithstanding the widespread clinical deployment of PD-L1 antibodies, their response rates remain approximately 20%, prompting investigations into alternative immunosuppressive pathways that may be implicated upon PD-L1/PD-1 axis inhibition[14, 15].

Sialic acids, comprising a class of negatively charged 9-carbon glycans linked to the termini of glycan chains via α-glycosidic bonds, serve as self-recognition signals on cell surfaces[16, 17]. Under normal physiological conditions, these signals transmit inhibitory cues to immune cells via Siglecs (Sialic acid-binding immunoglobulin-like lectins), establishing a state of self-tolerance and immune homeostasis[18]. However, pathogens or tumor cells exploit this mechanism, precipitating immune evasion and impeding effective clearance of cancer cells and pathogens by the immune system[19–21].

Our investigation unveiled a significant positive correlation between PD-L1 expression and genes associated with sialic acid in colorectal cancer. Treatment with a PD-L1 nanobody culminated in elevated cellular sialic acid levels, while sialidase treatment led to heightened cellular PD-L1 levels. These findings suggest that PD-L1 and sialic acid may represent two complementary immunosuppressive pathways. Consequently, we engineered a novel dual-function molecule by combining PD-L1 nanobody with sialidase protein (Nb16-Sia), which elicited potent in vivo anti-tumor effects. Subsequent immune deletion experiments underscored that Nb16-Sia primarily exerted anti-tumor effects through macrophages. Tumor-associated macrophages (TAMs), prevalent in various tumor types, are typically associated with a poor prognosis[22]. TAMs facilitate tumor growth and metastasis by suppressing immune responses and fostering a pro-tumor microenvironment[23, 24]. Our data revealed that Nb16-Sia could polarize macrophages to the M1 phenotype and reduce TAMs in tumor tissue.

## Materials and Methods

### Clinical data analysis

Clinical gene expression data of Colon adenocarcinoma (COAD) were obtained from the TCGA data sets (https://portal.gdc.cancer.gov/). The correlation between PD-L1 and other gene were analyzed by TIMER2.0 online tool and the gene expression difference in PD-L1 high and PD-L1 low group was analyzed by R software v4.0.3.

### Cell lines

CT26, MC38 and Raw264.7 cells were purchased from Procell. CT26 were cultured in RPMI-1640 medium (MA0215, Meilunbio) containing 1% P/S and 10% FBS. Raw264.7 cells were cultured in minimum essential medium (MEM; MA0217, Meilunbio) supplemented with 1% P/S and 10% FBS. HEK 293F cells were purchased from Thermo Fisher and grown in FreeStyle™ 293 expression medium (12338026, Thermo Fisher). All cells were cultured at 37°C in a 5% CO2 humidified atmosphere.

### Expression and purification of proteins

Genes encoding Nb16, Sialidase, and Nb16-Sia were cloned by GenScript Corporation into the pET28a vector for expression using BL21(DE3) cells. Above proteins were purified using Ni Sepharose™ 6 Fast Flow (175318, Cytiva). Endotoxins were removed using a high-capacity endotoxin removal spin kit (88275, Thermo Fisher). Gene encoding human Siglec9 (amino acid residues 18–347) was cloned into the pFUSE-hIgG1-Fc2 vector (InvivoGen) for expression using HEK293F cells. Siglec9-Fc protein was purified by Protein A Sepharose (17127901, Cytiva).

### Elisa assay for antibody binding activity

Elisa plates were coated with 100 μL of mouse-PD-L1 antibody (2 μg/mL) per well and incubated overnight at 4°C. Each well was blocked with 200 μL BSA (3%) for 1h at 37L. After washing, 100 μL of Nb16 and Nb16-SiaT antibodies were added to each well and incubated for 1 hour at 37L. Anti-His-HRP (dilution 1:10000, 30403, Yeasen) was added and incubated at 37L for 1 hour. After washing, 100 μl of TMB solution was added to each well and incubated at 37L for 10 minutes, followed by the addition of 50 μl of 2M H_2_SO_4_ to stop the reaction. The absorption value was measured at a 450 nm within 5 minutes.

### Flowcytometric and Immunophenotype Analysis

Raw264.7 cells were treated with Sialidase (75 μg/mL) or Nb16-Sia (100 μg/mL) for 24h. And then collected cells were incubated with Siglec9-Fc (2 μg/mL) in 100 μL volume. After washed with PBS, 1 μL of Anti-Human IgG FITC antibody was added and then the ready samples were detected by ACEA NovoCyte.

For immunotyping of intra-tumoral cells, tumor tissues were digested by collagenase IV (40510ES60, Yeasen) and hyaluronidase (20426ES60, Yeasen) and were filtered through 75-μm nylon mesh (7061011, Dakewe) into single-cell suspensions and then were subjected to cell extraction using lymphocyte separation medium (7211011, Dakewe) for sorting tumor-infiltrating lymphocytes (TIL) or were subjected to erythrocyte removal by red blood cell lysis buffer (40401ES60, Yeasen) for nonlymphocyte staining. Then, the cells were blocked with 4% FBS and anti-CD16/CD32 (553141, BD Biosciences), incubated with surface marker antibodies for 20 minutes at 4°C and then permeabilized with BD cytofix/cytoperm buffer (554714) before intracellular labeling antibodies were added for 30 minutes at 4°C. Then transcription factors were labeled for 45 minutes at 4°C by antibodies after being permeabilized with BD TF Fix/Perm buffer (562574). Flow cytometry analysis was performed using ACEA NovoCyte, and data processing was done through NovoExpress software (version 1.4.0). Antibody staining was performed following the manufacturer’s recommendations.

### Animal study

Female 6- to 8-week-old BALB/c and C57/B6 mice were purchased from the Shanghai Slack. Balb/c nu/nu mice were purchased from Vital River Laboratory. All mice were maintained under specific pathogen-free conditions in the animal facility of the Shanghai Institute of Materia Medica, Chinese Academy of Sciences (SIMM). Animal care and experiments were performed in accordance with protocols approved by the Institutional Laboratory Animal Care and Use Committee (IACUC).

For in vivo anti-tumor assay, CT26 or MC38 cells (1 × 10^6^) were injected subcutaneously into the right flank of Balb/c mice, C57/B6 or Balb/c-nu/nu mice. Mice were treated intraperitoneally or peri-tumor subcutaneously with Nb16 (25 μg), Sia (75 μg), or Nb16-Sia (100 μg) on day 5, day 7, day 9, day 11.

For macrophage deletion assay, macrophages were deleted using chlorophosphate-liposomes (200 μL/each/ i.p. 40337ES08, Yeasen) on days −2, 4, and 8. Endpoint dissected tumor tissues were made into paraffin sections and characterized by immunohistochemical staining for CD206 expression.

### RNA-Seq

Raw264.7 was treated with 100 μg/mL Sia or heated Sia for 24h, and then total RNA was extracted using Column Total RNA Purification Kit (B511361, Sangon). RNA transcriptome libraries construction, sequencing, and basic data analysis were conducted by Majorbio. Based on the RNA-seq raw data, differential expression was evaluated with DESeq. A fold change of 2:1 or greater and a false discovery rate (FDR)-corrected P value of 0.05 or less were set as the threshold for differential genes. GSEA was performed using GSEA software (http://www.broadinstitute.org/gsea).

### Western blot and Lectin blot

Proteins were detected using western blot analysis. Cells were washed with PBS and lysed in RIPA lysis buffer. Cell lysates were quantified for protein content using the BCA method. Subsequently, 10 µg protein/lane was resolved on SDS-PAGE (10% gel) and transferred onto a PVDF membrane. Membranes were blocked in 5% bovine serum album for 1h at room temperature. Subsequently, membranes were probed with primary antibodies against SYK (dilution 1:1000, 14858-1-AP; Proteintech), GAPDH (dilution 1:1000, A19056; Abclonal), iNOS (dilution 1:1000, A3774; Abclonal) Phospho-JNK (dilution 1:1000, AP1337; Abclonal) and JNK (dilution 1:1000, A0288; Abclonal) at 4°C overnight. Subsequent to washing, the membranes were incubated with secondary antibodies (dilution 1:5000, AS014; Abclonal) for 1 h at room temperature and visualized using the ECL Basic kit (RM00020, Abclonal). Glycans were determined by lectin blot. Different from the Western blot, biotinylated MAL-II lectin (10 μg/mL, B-1265-1; Vectorlab) was used for the primary antibody and Streptavidin-HRP (dilution 1:2000, A0303; Beyotime) for the secondary antibody.

### Statistical analysis

The in vivo experiments were randomized but the researchers were not blinded to allocation during experiments and outcome analysis. Statistical analysis was performed using GraphPad Prism 8 Software. A Student t test was used for comparison between the two groups. Multiple comparisons were performed using one-way ANOVA followed by Tukey multiple comparisons test or two-way ANOVA followed by the Tukey multiple comparisons test. Detailed statistical methods and sample sizes in the experiments are described in each figure legend. No statistical methods were used to predetermine the sample size. All statistical tests were two-sided, and P values < 0.05 were considered to be significant. Ns, not significant; *, P < 0.05; **, P < 0.01; ***, P < 0.001.

## Results

### Clinical database analysis revealed that PD-L1 and Sialic acid are co-related

Numerous enzymatic processes govern the transport of sialic acid within mammalian cells. Specifically, hydrolase and transferase enzymes play pivotal roles in modulating the abundance of sialic acid molecules residing on the cellular membrane[25]. Sialic acid transferase mediates the attachment of sialic acid moieties onto the cell surface, while sialic acid hydrolase facilitates their removal. Thus, the transcriptional regulation of these enzymatic components holds potential for reflecting the sialic acid profile at the cellular periphery. In this investigation, we elucidate a negative correlation between the expression levels of sialic acid hydrolases (NEU1, NEU4) and PD-L1, alongside a positive correlation between the expression levels of sialic acid transferases (ST3GAL1, ST3GAL5, ST3GAL6, ST6GALNAC5, ST8SIA1, ST8SIA4) and PD-L1 within colorectal cancer specimens sourced from the TCGA database (Fig. 1A). Furthermore, stratifying patients based on PD-L1 expression levels revealed significantly augmented sialic acid transferase expression and concomitantly diminished sialic acid hydrolase expression in individuals exhibiting heightened PD-L1 levels (Fig. 1B). To evaluate the interplay between sialic acid and PD-L1 expression, cellular models were subjected to PD-L1 antibody treatment, resulting in heightened sialic acid levels (Fig. 1C). Conversely, treatment with sialidase induced upregulated PD-L1 expression (Fig. 1D). These observations suggest a potential reciprocal modulation between the PD-L1 and sialic acid pathways. Consequently, we engineered nanobody fusion proteins possessing dual functionalities, namely PD-L1 blocking and sialidase activities, with the aim of concurrently targeting these immunosuppressive pathways.

**Figure1.**
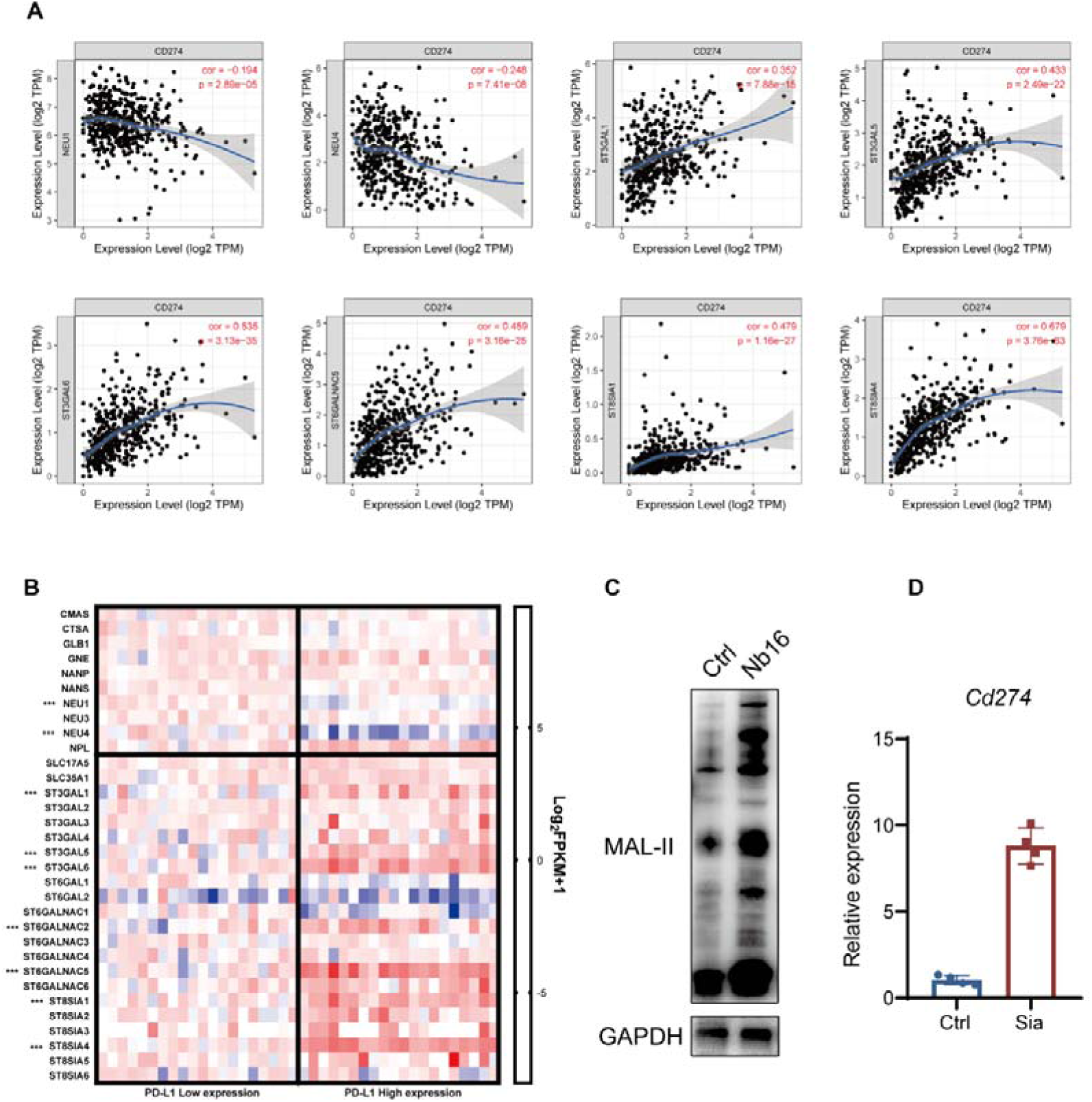
Correlation between PD-L1 level and sialic acid level in colorectal cancer tumor tissues. (A) Analysis of the correlation between key genes for sialic acid synthesis mRNA expression and CD274 (PD-L1) in tumors using the TIMER 2.0 tool. (B) Analysis of the fold change in key genes for sialic acid synthesis mRNA expression in the PD-L1 high group compared to the PD-L1 low group, compiled from TCGA data. (C) Lectin blot analysis of the sialic acid level in RAW264.7 cells after treated by PD-L1 nanobody (Nb16). (D) Quantitative PCR analysis of *Cd274* mRNA expression in RAW264.7 cells after treated by sialidase (Sia).

### Construction and validation of nanobody enzyme fusion protein

After analyzing the results above, a novel compound was synthesized by integrating PD-L1 nanobodies with sialidase. Prior investigations have evidenced the potent binding capability of PD-L1 nanobody within this composite, effectively inhibiting its functionality[26]. The sialidase component, sourced from the genome of *Actinomyces sp. oral taxon*, underwent cloning and subsequent validation of its enzymatic efficacy. A comprehensive series of assays revealed the remarkable thermal tolerance and heightened enzymatic performance of this sialidase (Fig. S1). Next, we conjugated PD-L1 nanobody and sialidase utilizing (G4S)3 linkers, followed by insertion into the pET28a vector (Fig. 2A). The obtained fusion protein was named as Nb16-Sia and successfully expressed, achieving a protein purity more than 90% (Fig. 2B). Subsequent assessment affirmed binding activity (EC_50_) towards PD-L1 of Nb16-Sia measured at 2.73 nM, consistent with the Nb16’s binding activity (Fig. 2C). Moreover, the enzymatic performance of Nb16-Sia mirrored that of Sia, as demonstrated in the pNP substrate assay (Fig. 2D). Furthermore, evidence was established confirming the enzymatic proficiency of Nb16-Sia in cleaving sialic acid residues on cellular surfaces, as indicated by Siglec-9 labeling (Fig. 2E-F). These collective findings mark the successful development of Nb16-Sia, a nanobody enzyme fusion protein endowed with dual functionalities, including PD-L1 binding and sialic acid hydrolysis.

**Figure2.**
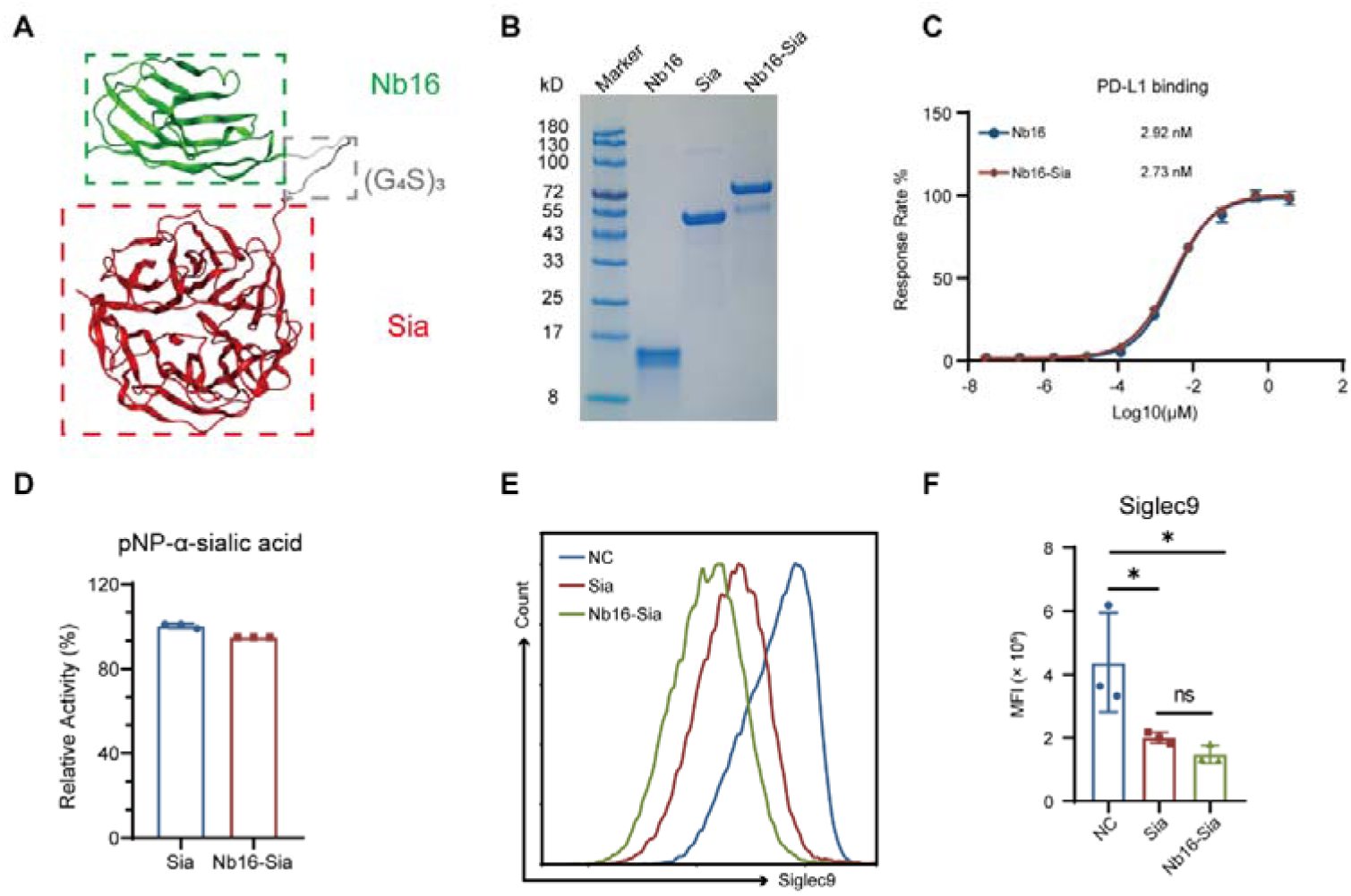
Construction and activity validation of the of Nb16-Sia. (A) The structure basis of the nanobody enzyme fusion protein Nb16-Sia predicted by AlphaFold2. (B) SDS-PAGE analysis of the obtained protein purified from *E. coli* using Caucasian blue stain. (C) Comparison of Nb16-Sia with Nb16 in PD-L1 binding affinity detected by ELISA. (D) Comparison of the sialidase cleavage activities of Sia and Nb16-Sia using pNP-α-sialic acid. (E) Flow cytometry detection of RAW264.7 cell surface sialic acid levels using Siglec9 labeling after enzymatic digestion of Nb16-Sia and Sia. (F) Quantitative analysis of median fluorescence intensity (MFI) in Fig. 2E.

### Nb16-Sia exerted anti-tumor effect in vivo and exhibited good safety

To evaluate the in vivo antitumor efficacy of Nb16-Sia, experiments were conducted using the CT26 tumor model. The treatment groups received four doses administered every two days, and therapeutic activity was compared across cohorts treated with PBS, Nb16, Sia, Combo, and Nb16-Sia. The results demonstrated a significant inhibition of tumor growth in mice treated with Nb16-Sia compared to those administered Nb16, Sia, and Combo (Fig. 3A-B). Notably, administration at a dosage of 5 mg/kg elicited a more pronounced and sustained antitumor response compared to the 2.5 mg/kg dosage, indicating a dose-dependent effect of Nb16-Sia on tumor suppression (Fig. 3C). To further validate Nb16-Sia’s antitumor effect, the in vivo model was expanded using MC38 tumor cells. As shown in Figure 3D, Nb16-Sia reduced tumor growth by 56%. Additionally, a comparison between peritumoral injection (PT) and intraperitoneal injection (IP) revealed that Sia (PT) more effectively suppressed tumor growth than Sia (IP), underscoring the necessity of the fusion of Nb16 with Sia. Collectively, these findings underscore the efficacious antitumor properties of antibody fusion proteins in vivo. Importantly, post-administration analysis of peripheral blood parameters revealed no significant deviations in body weight and liver and kidney function indicators across the five treatment groups (Fig. 3D-J), thereby reinforcing the safety profile of the drug candidate. Further assessment of the in vivo pharmacokinetic profile revealed that the half-life of Nb16-Sia is 6.32 ± 1.00 hours (Fig. 4A-B). The main PK parameters were shown in Table 1.

**Figure3.**
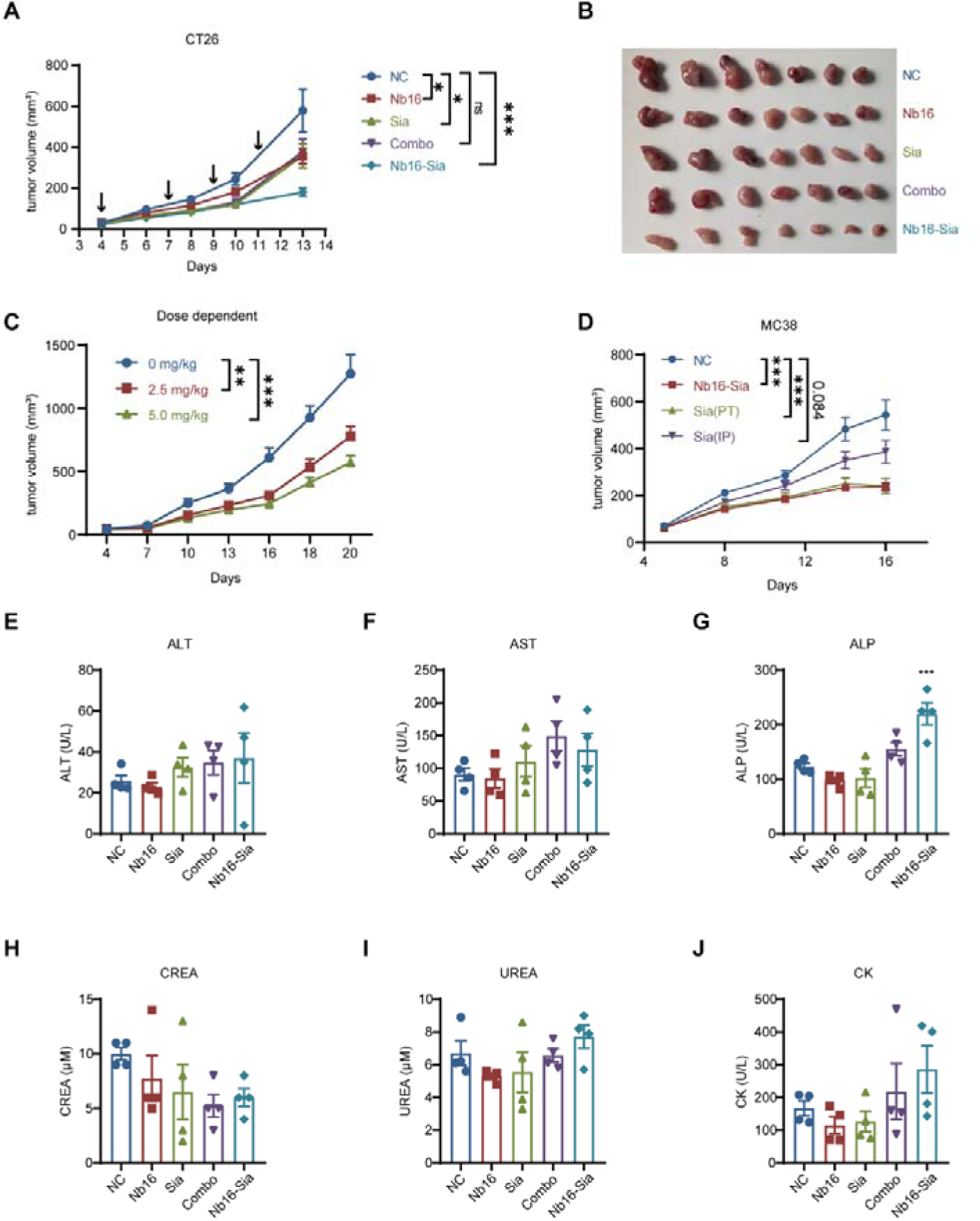
Anti-tumor effect and safety profile of Nb16-Sia in vivo. (A) Tumor volume growth curve in CT26 subcutaneous tumor model after receiving PBS, Nb16, Sia, Nb16-Sia or combination of Nb16 and Sia (n=7). (B) Picture of the subcutaneous tumor dissociated from sacrificed mice at the endpoint of the experiment. (C) Tumor volume growth curve in CT26 subcutaneous tumor model after administration of 2.5 mg/kg and 5 mg/kg dosages of Nb16-Sia. (D) Tumor volume growth curve in MC38 subcutaneous tumor model after receiving Sia (peritumor), Sia (intraperitoneal) and Nb16-Sia (n=8). (E-J) Liver (ALT, AST, ALP), kidney (CREA, UREA) and cardio (CK) function index among 5 groups after administration (n=4).

**Figure4.**
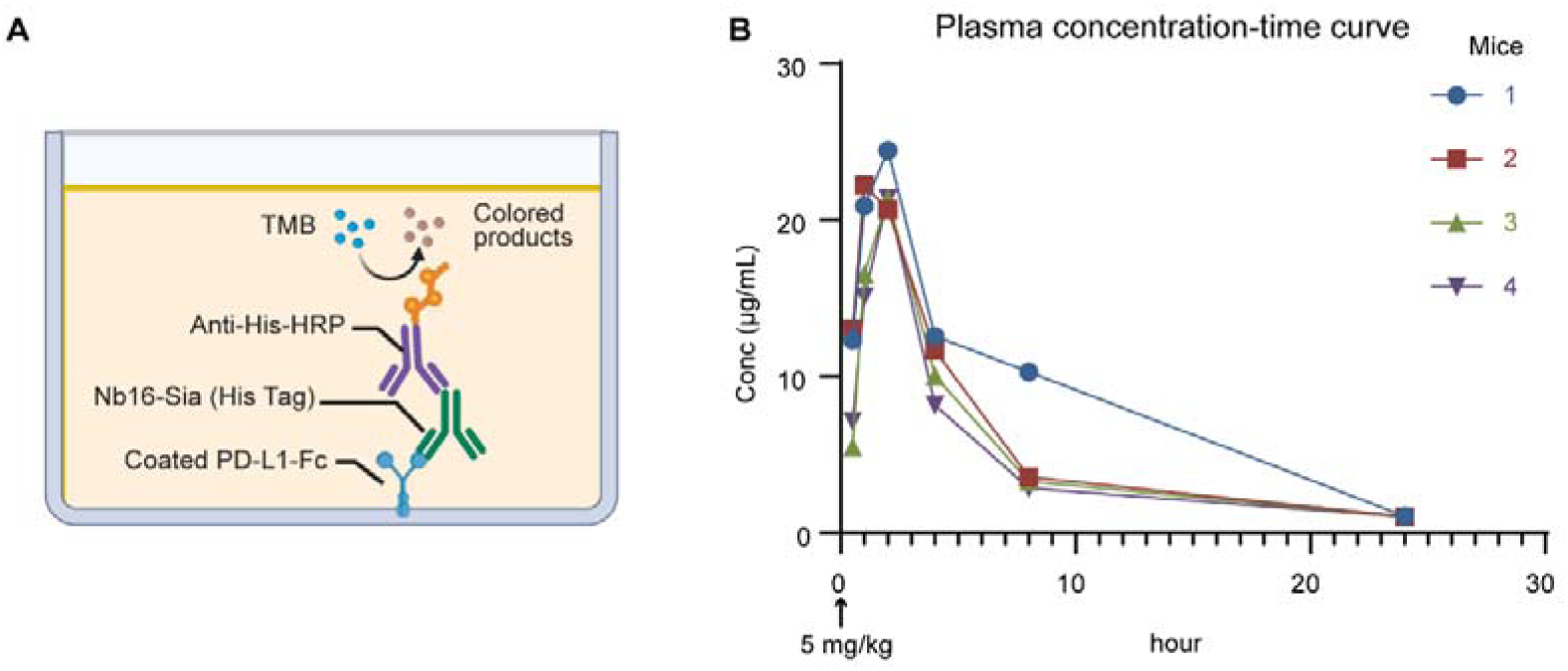
Concentration-time curve after intraperitoneal injection of a single dose of Nb16-Sia (5 mg/kg) in mice. (A) Schematic of detection assay in testing the blood concentration of Nb16-Sia. (B) Concentration of Nb16-Sia in mice plasma at 0.5h, 1h, 2h, 4h, 8h and 24h after injection. Each color represents an individual mouse.

**Table 1.** Pharmacokinetic parameters of Nb16-Sia after intraperitoneal injection of a single dose of Nb16-Sia (5 mg/kg) in mice.

| **animal** | **Tmax**<br>**(h)** | **Cmax**<br>**(ug/ml)** | **AUC<sub>all</sub>**<br>**(h\*ug/ml)** | **AUC<sub>(0-∞)</sub>**<br>**(h\*ug/ml)** | **Vd_F**<br>**(ml/kg)** | **Cl_F**<br>**(ml/h/kg)** | **MRTlast**<br>**(h)** | **T1/2**<br>**(h)** |
| --- | --- | --- | --- | --- | --- | --- | --- | --- |
| 1 | 2 | 24.5 | 208 | 217 | 181 | 23.1 | 6.14 | 5.45 |
| 2 | 1 | 22.3 | 134 | 142 | 281 | 35.2 | 5.33 | 5.54 |
| 3 | 2 | 21.4 | 119 | 129 | 379 | 38.8 | 5.56 | 6.77 |
| 4 | 2 | 21.4 | 109 | 120 | 452 | 41.6 | 5.57 | 7.52 |
| **Mean** | **1.75** | **22.40** | **142.50** | **152.00** | **323.25** | **34.68** | **5.65** | **6.32** |
| **SD** | **0.50** | **1.46** | **44.86** | **44.26** | **117.90** | **8.15** | **0.34** | **1.00** |

### The effect of Nb16-Sia mainly depended on macrophage

To further deepen our understanding of the immunological mechanisms underlying the action of Nb16-Sia, we conducted an analysis of the immune microenvironment within tumor tissues using flow cytometry. Our findings revealed a significant enhancement in the secretion of key cytokines such as IFN-γ, Perforin, Granzyme B, and TNF-α by T cells in response to Nb16-Sia (Fig. 5A), along with a mitigation of T cell exhaustion (Fig. 5B). Additionally, we observed a significant increase in NK cell infiltration within tumor tissues (Fig. 5C). In the Nb16-Sia group, there was an improvement in the expression of PD-L1 in macrophages, which corresponded to our data in Fig. 1D and could enhance the targeting of Nb16-Sia. Furthermore, the expression of MHC-II in macrophages was elevated, indicating an enhanced antigen-presenting capability (Fig. 5D). Collectively, these immunophenotyping results revealed an improved tumor immune microenvironment. Subsequent experimentation in T-cell-deficient murine models unveiled a diminished anti-tumor efficacy of Nb16-Sia, albeit with a persistent impediment on tumor growth (Fig. 5E), suggesting a partial reliance on T cells for its therapeutic activity. Within the tumor milieu, macrophages represent a substantial fraction of the infiltrating immune cell population alongside T cells[27]. Typically, macrophages undergo phenotypic alterations to assume a tumor-associated macrophage phenotype within the tumor microenvironment, thereby facilitating immune evasion[28]. We posited that macrophages might also exert a pivotal influence on the efficacy of Nb16-Sia. Indeed, experiments involving macrophage depletion resulted in a complete abrogation of the anti-tumor effects conferred by Nb16-Sia (Fig. 5F). Furthermore, immunohistochemical staining revealed a conspicuous reduction in the infiltration of CD206+ macrophages subsequent to Nb16-Sia treatment (Fig. 5G-H). In summation, our investigations identify Nb16-Sia as a potent inhibitor of tumor progression, achieved through the suppression of tumor-associated macrophage infiltration and augmentation of T cell functionality within the tumor microenvironment.

**Figure5.**
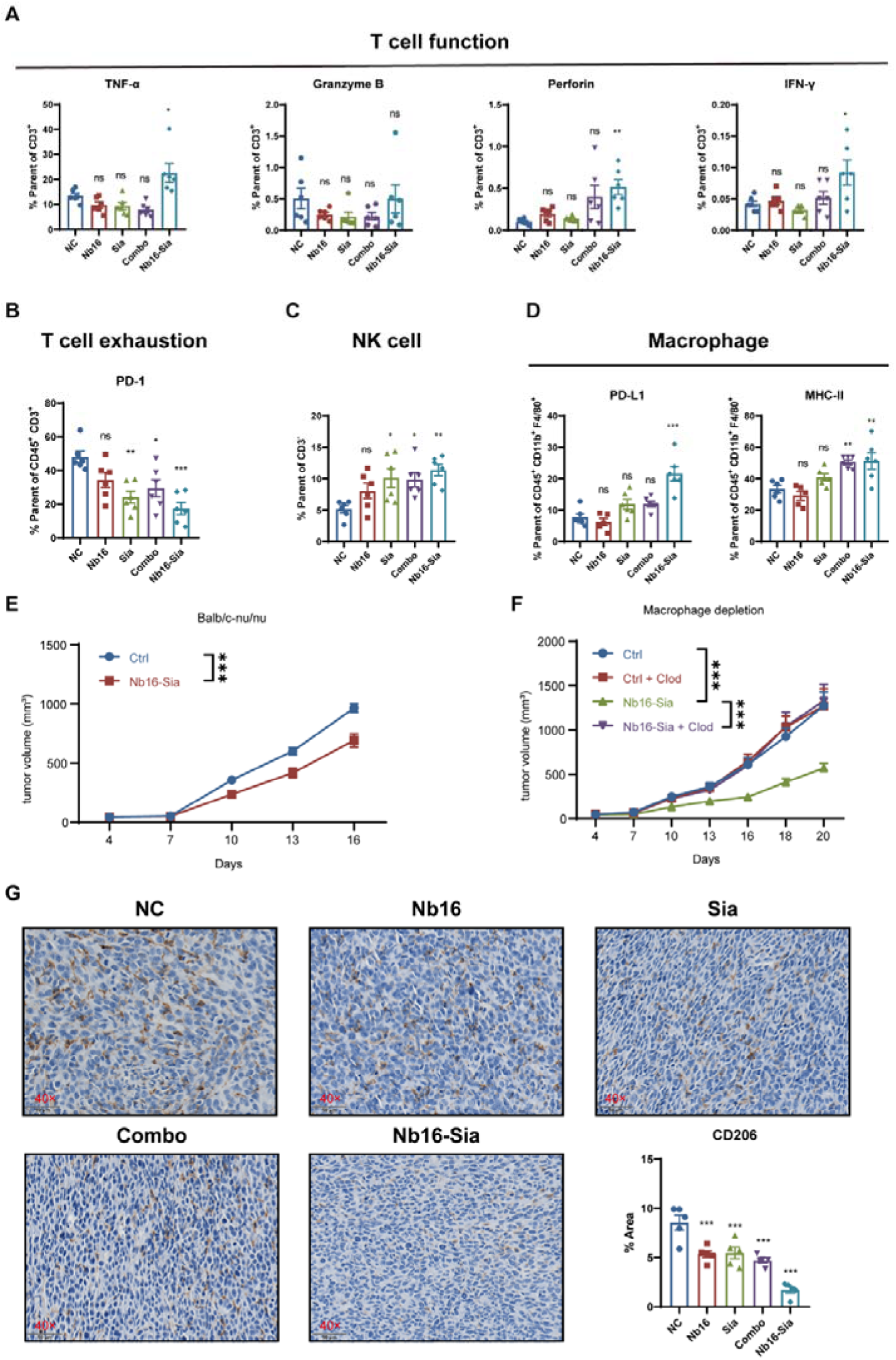
The reliance of Nb16-Sia anti-tumor effect on macrophages. (A-D) Immune effector cells in CT26 tumors from mice treated with 5 mg/kg of Nb16-Sia were quantified by flow cytometry (n = 6). (E) BALB/c-nu/nu mice bearing CT26 tumors were treated with Nb16-Sia (5 mg/kg). Tumor growth are shown (n=10). (F) Mice bearing CT26 tumors pre-treated with clodronate liposomes were injected with Nb16-Sia. Tumor growth is shown (n=9). (G) Representative images of IHC stain with CD206 in CT26 tumor tissues (40×).

### Nb16-Sia could promote M1 polarization by C-type lectin pathway

The aforementioned data demonstrates that Nb16-Sia exerts its anti-tumor effects primarily through modulation of macrophage activity. Despite extensive research elucidating the impact of the PD-L1 antibody on macrophages, our understanding of the influence of sialidase on macrophages remains incomplete. Thus, we conducted a transcriptomic analysis of macrophages post-sialidase treatment. The results unveiled a marked upregulation in the glycan-related C-type lectin pathway (Fig. 6A-B), accompanied by a notable elevation in SYK (spleen tyrosine kinase) expression within this pathway (Fig. 6C). Validation through Western blot analysis confirmed the enhancement of SYK levels by Nb16-Sia treatment, as well as activation of the downstream JNK (c-Jun N-terminal kinase 1) kinase pathway (Fig. 6D-E).

**Figure6.**
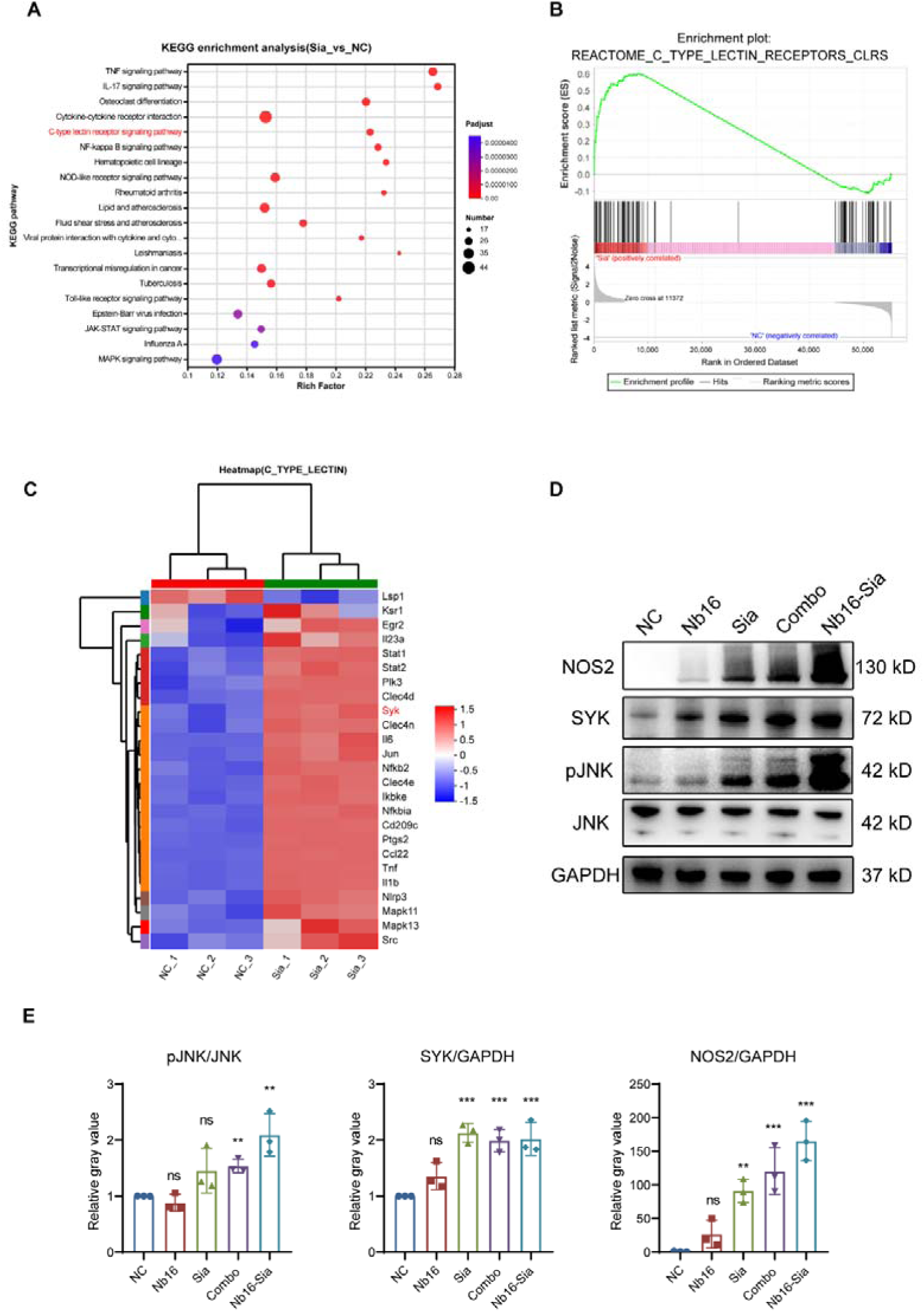
Nb16-Sia polarized macrophage to M1 phenotype through C-type lectin pathway. (A) KEGG analysis of the differential genes between Sia-treated group and negative control. (B) GSEA enrichment analysis of C-type lectin pathway. (C) Heatmap of the differential genes in C-type lectin pathway. (D) Representative image of western blot in detecting SYK, JNK and NOS2 expression at protein level. The quantitative results are analyzed by gray value and shown (E). Table1 Pharmacokinetic parameters of Nb16-Sia after intraperitoneal injection of a single dose of Nb16-Sia (5 mg/kg) in mice.

## Discussion

In this investigation, a novel fusion protein comprised of PD-L1 nanobody and sialidase was developed and designated as Nb16-Sia. In vivo assessments revealed that Nb16-Sia elicited anti-tumor effects by modulating macrophage activity. In vitro experiments showed that Nb16-Sia could induce activation of the C-type lectin pathway in macrophages, thereby promoting polarization toward the M1 phenotype. Antibody fusion proteins have garnered increasing attention as a promising avenue for bispecific antibody development. When formulating the molecular structure of antibody fusion proteins, it is imperative to consider both the intended target and the architectural configuration of the antibody, alongside the rationality of its fusion with the coupling agent[29]. PD-L1 exhibits high expression levels in both tumor cells and TAMs[30, 31], with a notable correlation observed between sialic acid levels and PD-L1 expression in tumor tissues (Fig. 1A). Consequently, we elected to merge PD-L1 with sialidase and deliver the resultant fusion protein to tumor tissues utilizing PD-L1 antibodies. In the realm of cancer therapy, nanobodies present significant advantages in terms of tumor targeting and rapid clearance, thereby facilitating optimal delivery to tumor cells while minimizing adverse effects on healthy tissues[32]. Furthermore, nanobodies offer facile modification, hence PD-L1 nanobodies were selected as the focal targets in this investigation. Given that sialidase from mammalian cells typically localizes within lysosomes[33, 34], its enzymatic activity is primarily exerted under acidic conditions[35]. Accordingly, sialidase was sourced from oral commensal bacteria due to its superior pH tolerance range. This study represents the inaugural endeavor in the field of PD-L1/sialic acid dual-targeting nanobody enzyme fusion proteins.

Tumor-associated macrophages (TAMs) represent a distinct subset of macrophages that infiltrate neoplastic tissues, primarily originating from circulating monocytes[36, 37]. The recruitment of monocytes from the circulation into the tumor microenvironment (TME) is facilitated by chemotactic factors such as CSF1 and CCL2 secreted by tumor cells[38–40]. Upon reaching the TME, monocytes undergo differentiation into macrophages. Initially perceived to possess anti-tumorigenic properties, TAMs are now acknowledged to exhibit pro-tumorigenic and immunosuppressive characteristics[41]. TAMs contribute to the progression of tumors through various mechanisms, including the promotion of tumor cell proliferation, facilitation of invasion and metastasis, facilitation of immune evasion, and induction of angiogenesis[42–45]. Furthermore, TAMs play a regulatory role in tumor cell metabolism, notably by fostering aerobic glycolysis and malignant progression[46]. Notably, studies have indicated that TAMs constitute approximately 20% of tumor tissue and manifest heightened expression of PD-L1. This suggests a potential mechanism underlying the efficacy of PD-L1 antibody therapies in tumor patients, possibly through the targeting of macrophages. Henceforth, we hold that TAMs play the role of “umbrella” for tumor cells, and targeting TAMs can be a promising approach for anti-tumor therapy[47–49].

However, it is imperative to acknowledge the limitations inherent in our study, necessitating further investigation. Firstly, there is a crucial need to explore the potential systemic toxicity of sialidase on circulating monocytes subsequent to intravenous administration. Secondly, the immunogenicity of sialidase warrants thorough consideration[50]. Given that the production of anti-drug antibodies could impact the long-term safety profile of the drug, further efforts towards its humanization are warranted. Thirdly, the necessity for fusion of the Fc segment remains under deliberation due to concerns regarding the short in vivo half-life of Nb16-Sia, which could potentially compromise its efficacy[51]. Lastly, comprehensive studies are warranted to elucidate the precise role of Nb16-Sia in tumor progression and to identify adjunctive therapeutic agents capable of augmenting its antitumor efficacy.

## Notes

### Competing Interest Statement

The authors have declared no competing interest.

### Summary of Updates

The author order updated, and some minor error corrected in figure 5 legend.

