## Supplemental Figure 1 for "A nanobody-enzyme fusion protein targeting PD-L1 and sialic acid exerts anti-tumor effects by affecting tumor associated macrophages"

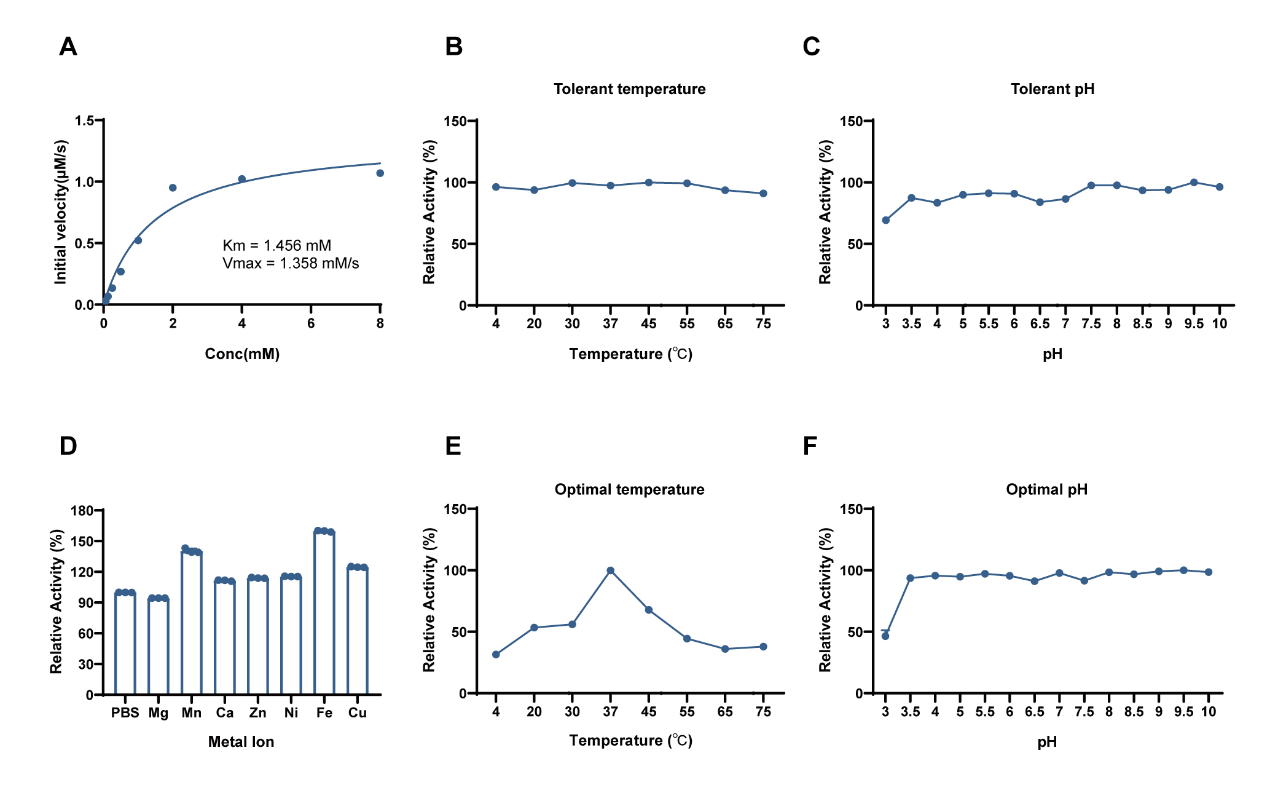

**Figure S1. Stability of Sialidase.** (A) enzyme kinetics curve of sialidase. (B) The tolerant temperature of Sialidase. (C) The tolerant pH of Sialidase. (D) Effect of variety of ions on Sialidase activity. (E) The optimal temperature of Sialidase. (F) The optimal pH of Sialidase.
